# Deletion of an sRNA primes development in a multicellular bacterium

**DOI:** 10.1101/2024.05.04.592516

**Authors:** Marco La Fortezza, Jasper Verwilt, Sarah Cossey, Sabrina Eisner, Gregory J. Velicer, Yuen-Tsu N. Yu

## Abstract

Small non-coding RNAs (sRNAs) are essential in regulating gene expression during many biological processes. The myxobacteria gene *pxr* encodes an sRNA known to block fruiting-body development, an aggregative multicellular process triggered by starvation. Deletion of *pxr* allows *Myxococcus xanthus* cells to develop in the presence of nutrients. However, potential Pxr binding targets and most genes regulated by Pxr remain unknown. Here, we found that the absence of *pxr* expression dramatically alters the temporal dynamics of development, thus suggesting an important new role of this sRNA in myxobacterial ecology. We transcriptionally profiled vegetative cells of *M. xanthus* strains possessing vs lacking *pxr* and found that over half of the genes impacted by *pxr* deletion during growth are linked to development, including known and potentially novel critical regulators. Many other genes are associated with general metabolic processes, which Pxr regulates positively. Our study discovers new phenotypic effects of Pxr regulation of likely ecological importance, identifies the suite of genes this sRNA controls during vegetative growth, reveals a previously unknown developmental regulator and provides new insights into the early molecular regulation of myxobacterial development.

## INTRODUCTION

Small non-coding RNAs (sRNAs) play an important role in regulating gene expression in bacteria, and their activity is essential for a wide variety of biological processes^1–3^. These include forms of virulence, stress responses, quorum sensing, and the initiation of aggregative multicellular development in the myxobacteria^3–5^. sRNAs regulate their targets through various molecular mechanisms, such as altering post-transcriptional transcript abundance or directly interfering with ribosome binding to impact translation^2,3^. sRNAs are distinguished as *cis-* or *trans-*acting relative to localisation of their gene targets^1,4,6^. The study of sRNAs is important for understanding the biological processes with which they are associated and their evolution.

Upon starvation, myxobacteria cells respond by aggregating and cooperatively developing into multicellular spore-bearing fruiting bodies^7^. The stringent response initiates fruiting-body development through a complex network of molecular interactions triggered when scarcity of amino acids impedes protein synthesis^8^. The molecular factors participating in fruiting-body formation are only partially known, although substantial progress has been made in recent years^8–11^. One major regulator of early development in the model species *Myxococcus xanthus* is the sRNA Pxr, which guards the transition from vegetative growth to multicellular development^5^.

The gene encoding Pxr (*pxr*) was discovered in a strain named Phoenix (PX) that spontaneously re-evolved the ability to make fruiting bodies and spores after its ancestral lineage had lost that ability^12^. A single-nucleotide substitution in PX localised in the *pxr* gene inactivates the sRNA function, restoring developmental proficiency relative to the developmentally defective strain OC from which PX derived^12^. The gene *pxr* encodes for the so-called long Pxr (Pxr-L) sRNA that, in vegetative cells, is subsequently cleaved into a shorter version (Pxr-S). Pxr-S levels decrease as soon as cells sense starvation, whereas Pxr-L levels remain stable throughout development^5^. A recent study revealed the existence of a much larger Pxr precursor transcript (Pxr-XL), and maturation to Pxr-S involves at least a two-step process regulated by the housekeeping ribonuclease RNaseD^13^. We previously hypothesised that Pxr-S is the key inhibitor that arrests aggregate formation and, thus, the entire process of fruiting-body development (Fig. 1a)^5^. However, whether Pxr-L directly influences fruiting-body formation is still unknown^13^.

**Figure 1.**
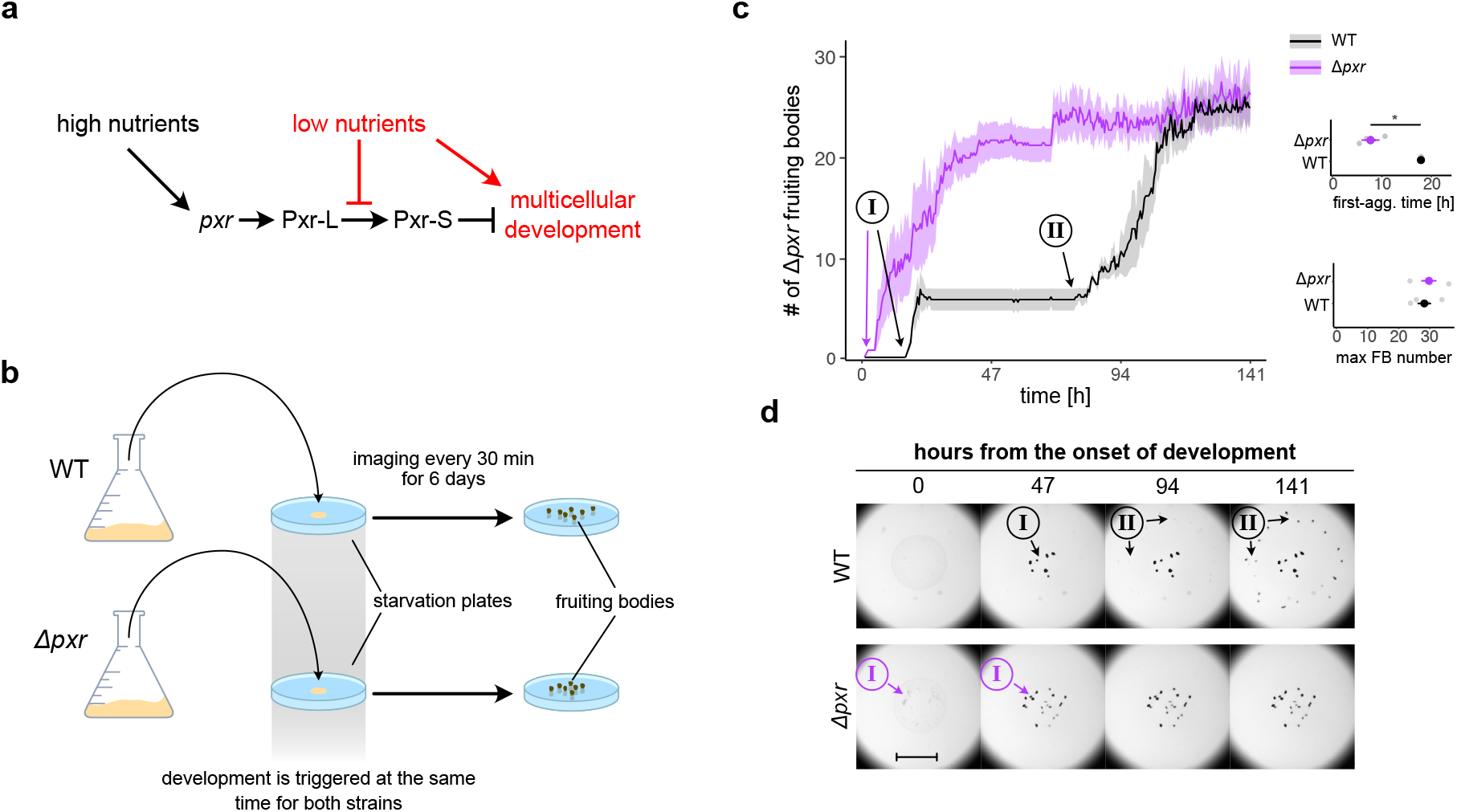
Deletion of *pxr* accelerates fruiting-body development during starvation. a) Simple model summarising the current knowledge of nutrient-level effects on Pxr processing and inhibition of multicellular development. **b)** Overview of the experiment used to compare the dynamics of fruiting-body development between WT and *Δpxr*. WT (GJV1) and mutant cells were independently grown in reach media until the mid-log phase and plated at equal density on starvation plates (minimal media) to induce multicellular development. Plates were imaged for 141 h (∼ 6 days) every 30 min to quantify the speed and proficiency of fruiting-body development over time. **c)** Plot reporting the average number of fruiting-body counts (thick lines) over time for WT (GJV1) (black line and shaded area) and *Δpxr* cells (purple line and shaded area). The two developmental waves observed in GJV1 are indicated with I and II circled in the plot area (black arrows); the first and only major developmental wave observed for *Δpxr* is also indicated with I and a purple arrow. Graphs on the right report the average developmental time at which aggregates first formed (top) and the average number of fruiting bodies present at the end of the experiment (bottom) for WT and *Δpxr* cells (black and purple, respectively). The shaded areas in the large plot and the error bars in both graphs represent the standard errors associated with each measurement (n = 4). The asterisk indicates a significant mean difference. The experiment was run using the same experimental protocol shown in Fig. 3a. **d)** Representative images of developing fruiting bodies (dark spots) over the time course of WT (GJV1) (top row) and *Δpxr* cells (bottom row). Arrows indicate the groups of fruiting bodies characterising the two developmental waves I and II circled in the images for both WT (black arrows and text) and *Δpxr* cells (purple arrows and text). Images reporting *Δpxr* developing cells lack the second developmental wave II present for the WT cells. The scale bar equals 1 mm.

Starvation-induced fruiting-body formation is present in most myxobacterial species identified to date^7,14^. Interestingly, homologs of the *M. xanthus pxr* variant are present in multiple species belonging to the suborder Cystobacterineae, within which *pxr* emerged^15^. When inserted into a deletion-mutant of *M. xanthus* lacking *pxr*, these inter-specific *pxr* homologs can partially or fully restore the developmental phenotype associated with the native *pxr* gene, thus suggesting a conserved function of Pxr across myxobacterial species^16^. While the development-inhibiting function of Pxr is known and plausibly similar in all species that express this sRNA, both the direct binding targets of Pxr and the set of genes ultimately regulated by Pxr in any species remain uncharacterised.

Previous studies have shown that Pxr is a developmental gatekeeper that blocks the transition from vegetative growth to fruiting-body formation when nutrients are still available^5^. Therefore, the absence of Pxr facilitates the activation of multicellular development even when nutrient levels remain high enough to fuel robust vegetative growth^5^. This result implies that the expression of some genes necessary for progression through early development is not regulated directly by nutrient level per se but rather by the active form of Pxr (Pxr-S), the level of which is regulated by external nutrient levels (by a still-unknown mechanism). Because Pxr-S is present at high levels during vegetative growth^5^, its absence due to deletion of *pxr* is expected to result in increased expression of some developmental genes already during growth. A mutant lacking *pxr* is thus likely to both: i) exhibit earlier progression to development upon starvation than its parent in which *pxr* is intact, and ii) even during vegetative growth, detectably express known positive regulators of development that are silenced during growth when *pxr* is intact.

Our data shows for the first time the molecular targets and effects of Pxr sRNA on development dynamics, which could significantly affect *M. xanthus* ecology and multicellular evolution.

## RESULTS

### The absence of pxr expression alters the temporal dynamics of fruiting-body formation

Impaired expression or functionality of *pxr* was previously associated with activation of the *M. xanthus* developmental program even under high-nutrient conditions that normally block development^5,12^ (Fig. 1a). Motivated by previous informal observations, we asked whether the absence of *pxr* expression during vegetative growth might accelerate development upon vegetative cells being placed in a development-inducing environment^5^. Thus, we initiated development on buffered agar and monitored the dynamics of fruiting-body formation by WT cells and cells devoid of *pxr* (GJV1*Δpxr*, also referred to hereafter as *Δpxr,* see Methods*: Strains, culturing conditions and induction of development*) for 141 hours by microscopy (ca. six days) (Fig. 1b) (see Methods*: Image acquisition and analysis*). As expected, both strains formed fruiting bodies when starving, but their development dynamics differed significantly (two-way ANOVA, time:strain, F = 500, *p* < 0.001) (Fig. 1c and d). *Δpxr* cells formed visible aggregates earlier (Wilcoxon test relative to the onset of aggregate formation, W = 16, *p* = 0.028) and reached their maximum fruiting-body number faster than WT cells (Wilcoxon test comparing the time point at which the maximum number of fruiting bodies formed, W = 0, *p* = 0.029) (Fig. 1c and d).

Notably, WT cells reached their maximum number of fruiting bodies (under our experimental conditions) through two waves of development (I and II in Fig. 1c and d). The first wave emerged near the plate centre (at the inoculation site) after ∼24 hrs. In contrast, the second wave formed an outer ring of fruiting bodies ∼50 hrs later (Fig. 1c). Similar developmental dynamics are not uncommon. Concentric rings of fruiting bodies can frequently be observed during development progression (unpublished observations). Interestingly, the second developmental wave characteristic of WT cells was absent in *Δpxr* cells, which formed most fruiting bodies within the first two days and reached nearly the maximum number within three days (Fig. 1c and d). Notably, the total number of fruiting bodies produced by the two strains at the end of the experiment was similar (Wilcoxon test relative to the last time point, W = 6, *p* = 0.6612) (Fig. 1c).

Fruiting-body formation is a developmental process that relies on quorum sensing and, thus, on cellular density^17^. Hence, we asked whether the observed differences in the dynamics of development of *Δpxr* mutant could be partially explained by altered sensitivity to cell density. However, the fruiting-body number for the *Δpxr* mutant cells in the presence and absence of nutrients was proportional to the initial inoculum density (Supp. Fig. 1a and 1b, respectively), and similar to WT when developing on buffered agar (Supp. Fig. 1b)^17^. Thus, no apparent difference in sensitivity to density on buffered agar was observed between the two strains (Supp. Fig. 1b).

Our analysis of developmental dynamics shows that lack of *pxr* expression during growth in liquid induced early activation and acceleration of the developmental program.

### Pxr positively regulates many metabolism genes during growth

Little is known about the genetic interactions of *pxr,* despite its important role during myxobacterial development. We used RNA-seq to compare genome-wide levels of gene transcripts during vegetative growth between *Δpxr vs* WT cells (Supp. Fig. 2a and b, see Methods: *Strain and culturing conditions*). Overall, the normalised read counts per gene between the WT and *Δpxr* profiles were highly correlated (Pearson’s *r* = 0.814) (Fig. 2a), as were counts between individual experimental replicates for both genotypes (Supp. Fig. 3a and b). Yet, the lack of *pxr* expression altered the transcript abundance of 299 genes (∼4% of all DK1622 annotated genes) relative to the wild type during vegetative growth (Supp. Table 1). Of those, 49.2% showed a significant decrease, and 50.8% showed an increase in the transcript levels due to the deletion of *pxr* (Supp. Fig. 2c) (see Methods: *RNA extraction and RNA-seq analysis*).

**Figure 2.**
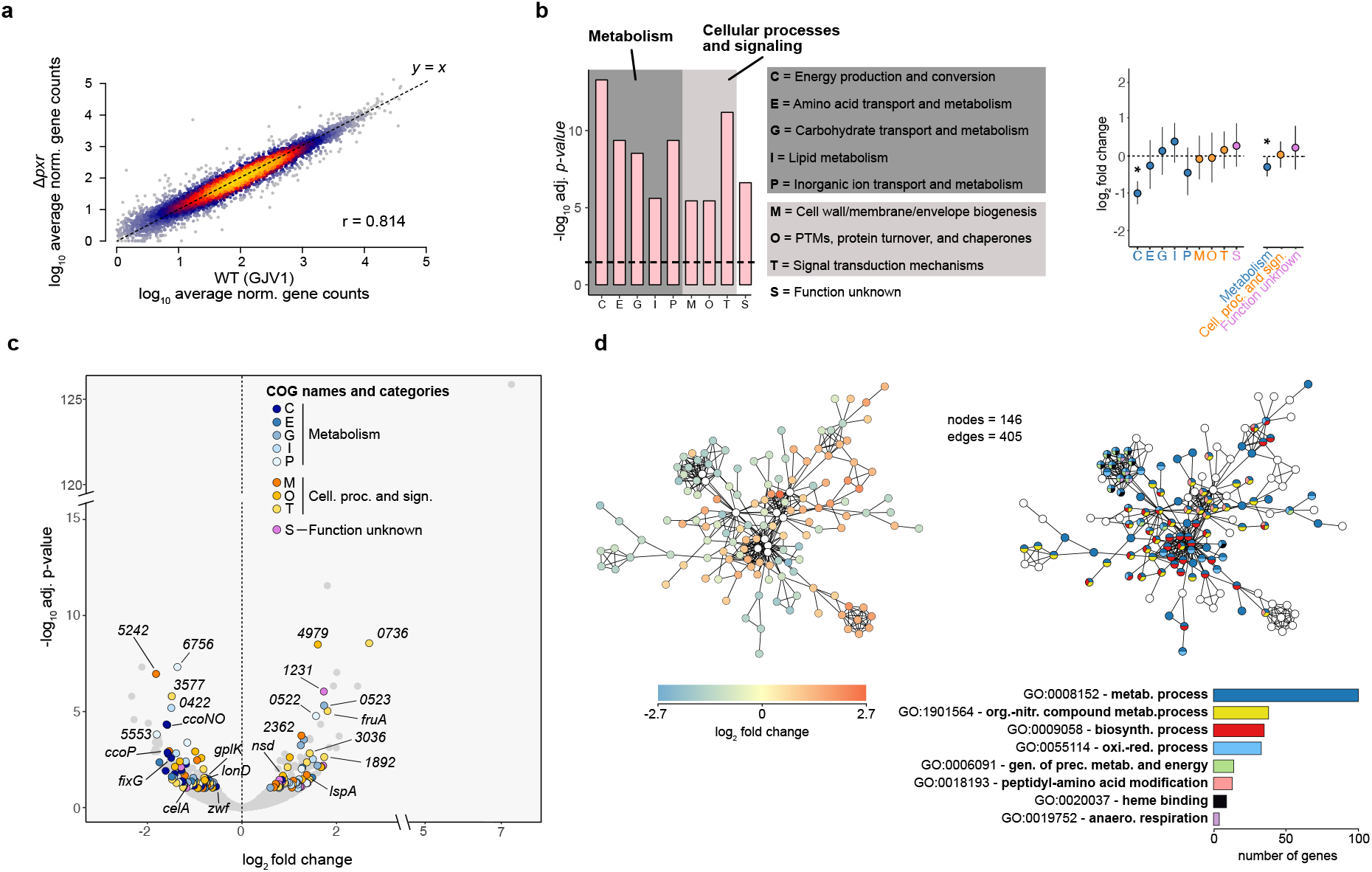
*pxr* expression downregulates many metabolism genes and upregulates many genes associated with cellular processes and signalling. a) Heat-density scatter plot reporting the average read counts per gene in both WT (GJV1) and the Δ*pxr* mutant. The colour gradient shifting from blue to yellow indicates increased data-point density. Pearson’s correlation *r* is shown within the plot area. **b)** Letf: COG categories significantly enriched with differentially expressed (DE) genes (Table S2). The *x* axis shows the enriched COG categories, while the *y* axis reports the significance levels expressed as -log10. The dashed black line highlights the significance threshold of *p* = 0.05. Dark and light grey areas group COG terms that belong to “Metabolism” and “Cellular processes and signalling”, respectively. Right: Dot plots reporting the average gene expression level grouped by enriched COG term (left) and by enriched COG category (right). In all cases, blue shaded dots refer to COGs terms associated with *Metabolism* (C, E, G, I, P), orange with *Cellular Processes and Signalling* (M, O, T), and purple with *Function unknown* (S). For both graphs, error bars represent a bootstrap 95% CI. **c)** Volcano plot reporting differential gene expression levels as a function of their adjusted p-values. Coloured dots refer to genes associated with enriched COGs categories (see panel **c**), while grey dots indicate genes that are either not significant (*adjusted p* > 0.1) or not annotated to any enriched COG term. Numbers refer to MXAN gene names. For the complete list of DE genes and their COG annotation, refer to Supp. Table S1. Visit this <u>link</u> to explore the graph in detail. **d)** Structure of the primary gene-interaction network found among the DE genes. Each node represents one gene, while each connecting line (edge) indicates the potential interaction between two genes. For the left version of the network, the colour gradient from blue to red indicates differential gene-expression levels between WT and Δ*pxr*. White dots represent genes that did not show a significant difference in expression between the two genotypes added to complete the interaction network. For the right version of the network, each colour represents one enriched GO term found among genes constituting the shown network (Supp. Table 3). Multicolour dots indicate genes annotated to multiple enriched GO terms. Grey dots represent genes not belonging to any enriched GO term. The bar plot below summarises the gene counts per GO term. The depicted networks can be downloaded and examined in more detail online using this **<u>link</u>**.

We found that 33 operons were significantly enriched for the presence of differentially expressed genes. Only one of these operons contained genes that changed expression discordantly – both up and down (operon ID = 0521: *MXAN_0975*, *MXAN_0976*, Supp. Fig. 3d, Supp. Table 2). All other operons’ genes changed transcriptional levels concordantly – up or down (two-tailed χ^2^ (1, *N* = 33) = 29.121; *p* = 0.0001) (Supp. Fig. 3d, Supp. Table 2). The overall directional consistency of transcriptional change among genes belonging to the same operon suggests an upstream function of Pxr to the expression or stability of the polycistronic transcripts encoded by these operons.

We then determined clusters of orthologous genes (COGs) (see Methods: *RNA extraction and RNA-seq analysis*) enriched among the differentially expressed genes. Nine COG categories were significantly enriched, although only a few genes constituted each category (Fig. 2b and c, Supp. Table 3). Five of the enriched categories were representative of *Metabolic processes* (categories: C, E, G, I, and P), three are related to *Cellular processes and signalling* (categories: M, O, and T), and one (S) contains genes related to the class *Function unknown*. Notably, most genes with decreased expression levels belonged to those COGs associated with metabolism (Fig. 2c,d). In contrast, genes with unaltered or increased expression were more likely to belong to categories related to cellular processes and signalling (Fisher’s exact test, *p* = 0.0105) (Fig. 2b,c Supp. Table 3).

We then extracted predicted protein interactions among the products of the differentially expressed genes from the STRING database^18^ (see Methods: *RNA extraction and RNA-seq analysis*). Our analysis individuated one distinct major network consisting of 405 total potential interactions among a subset of 145 genes (respectively the number of edges and nodes in the network) (Fig. 2d, and Supp. Fig. 4a). All of the other differentially expressed genes belonged to either minimal networks (2 or 3 nodes with 1 or 2 edges, respectively) or remained unmatched given their current annotation (Supp. Fig. 4a, Supp. Network Images). Interestingly, when qualitatively considering the distribution of transcriptional levels mapped onto the large network, genes with a similar deviation in their expression values are frequently associated with one another, again suggesting a potential role of Pxr acting upstream to these genes’ transcription (Fig. 2d, Supp. Fig. 4b).

A follow-up analysis of enriched gene ontology (GO) terms on this network revealed that a large proportion of genes were associated with metabolism (Fig. 2d, Supp. Table 4). Among them, for example, we identified genes involved in key metabolic pathways such as the genes *glpK, celA, zwf,* which are known to be respectively associated with the metabolism of glycogen, sucrose, and the pentose phosphate pathway^19–23^. We also identified *ccoNO*, *ccoP*, and *fixG,* which encode for cytochrome-associated proteins whose function is involved in the oxidative phosphorylation^24^ (Fig. 2c and Supp. Fig. 4b), as well as genes associated with the translational process, such as the two genes *fusA* (*MXAN_4082*) and *alaS*, respectively encoding for the ribosome elongation factor G and mediator of the alanyl-tRNA biosynthesis (see Supp. Table 1 and 3 for more details on the annotations for the individual functions of all differentially expressed genes)^25,26^.

We additionally generated a user-friendly online application using Shiny with R software (https://jzego7-marco.shinyapps.io/rnaseq_app/) to interactively visualise the presented data more thoroughly (see Methods: *RNA extraction and RNA-seq analysis* and Supp. Code).

Our global analysis of differentially expressed genes indicates that the absence of *pxr* expression during vegetative growth influences the transcript levels of a subset of genes linked to metabolic processes and cellular signalling, which tend to be positively and negatively regulated by Pxr, respectively (Supp. Tables 1, 3 and 4).

### Most genes affected during growth by deletion of pxr are associated with fruiting-body development

Given the role of *pxr* expression in controlling fruiting-body development and its dynamics, we sought to identify known developmental genes whose transcriptional levels during growth were altered by deletion of *pxr*. Hence, we compared our RNA-seq data to the most recent lists of genes associated with fruiting-body development in *M. xanthus.* Thus, we extracted and combined lists of developmental genes from the independently published works of Muñoz-Dorado et al.^9^, Sharma G. et al.^10^, and McLoon et al.^11^.

We first compared the degree of similarity between all these studies and ours. We contrasted the gene-transcript levels scaled around their relative average value for the WT cells during vegetative growth (Supp. Fig. 5a) (see Methods: *Comparison of RNA-seq profiles*). This comparison allowed us to estimate the consistency across the RNA-seq protocols used in all the studies based on the only physiological and experimental conditions they had in common, including ours. From these comparisons, we found that the vegetative profiles of WT cells for all three studies correlated with ours similarly (Supp. Fig. 5a). We thus retained all three studies for further comparisons and indicated which paper(s) associates a given gene with fruiting-body development (Supp. Table 5).

Notably, these three studies explored the expression details of potential developmental genes beyond our study’s scope (*i.e.*, not only during vegetative growth but also over periods of starvation). However, from all three studies, we determined the list of developmental genes by considering any gene found to change expression during starvation relative to their expression during vegetative growth. While ∼21% of all *M. xanthus* genes have been associated with development by any of the three development- transcriptome studies, ∼67.6% of genes differentially expressed between WT and *Δpxr* are found among those development-associated genes. Thus, gene regulation by Pxr is biased toward development-associated genes (binomial test, *p* < 10^-16^).

Of the developmental genes found to be differentially expressed in *Δpxr*, 64.4% were associated with development by Sharma et al.^10^, 59.9 % by McLoon et al.^11^, and 53.5% by Muñoz-Dorado et al.^9^(Fig. 3a and b, Supp. Fig. 5b). In addition, the so-obtained developmental genes constituted half of the large interaction network shown in Fig. 2d (non-developmental genes = 48%; developmental genes = 52%, Supp. Fig. 6) and 75% of the previously identified 33 enriched operons contained genes associated with development (Supp. Table 2 and Supp. Fig. 3d). Of the 311 genes related to development by all three transcriptome studies, only 42 (13.5%) were differentially expressed in the *Δpxr* mutant (Fig. 3a and c). Interestingly, a significant majority of these 42 genes showed increased transcriptional levels in *Δpxr* cells (30/42; Fisher’s exact test, *p* = 0.0028) (Fig. 3c). Among them, we identified well-known regulators of development, such as *fruA, nsd,* and *MXAN_0736,* which are, respectively, a crucial transcriptional regulator of other developmental genes, a nutrient-depletion sensing factor, and a histidine protein kinase relevant to spore germination^11,27–29^. We confirmed that the increased transcriptional levels of *fruA* were indeed reflective of increased protein abundance, with FruA being detectable already during vegetative growth in *Δpxr* mutant cells, whereas it became detectable only after six hours of starvation in WT cells (Fig. 3d). Other known regulators of development identified by only one or two of the development-transcriptome studies were also found to be differentially abundant by deletion of *pxr*, for example, *abcA*^30^*, lonD*^31^*, sgmE*^32^*, oar*^33,34^, and *rodK*^29,35^ (Fig. 3b, Supp. Table 1 and 5). Together, these data identify many developmental genes that Pxr regulates during vegetative growth, whether directly or indirectly.

**Figure 3.**
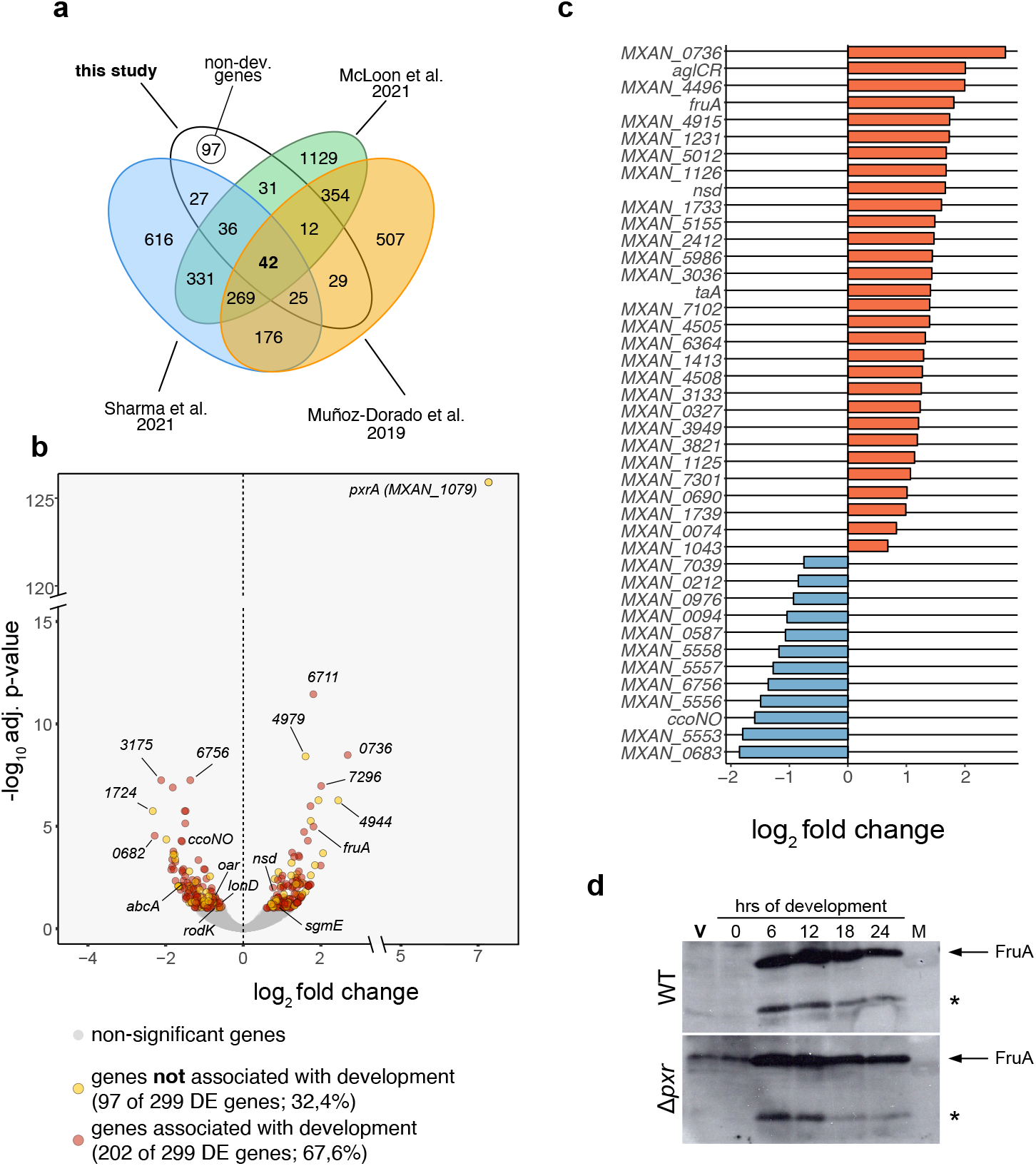
Lack of *pxr* changes the expression of many genes associated with fruiting-body development during vegetative growth. a) Venn diagram showing the number of DE genes found in our study (white area) that were identified as showing development-specific patterns of gene expression by Muñoz-Dorado et al. 2019^9^ (orange area), Sharma et al. 2021^10^ (blue area), or McLoon et al. 2021^11^(green area). (For details on the comparison, consult Methods: *RNA-seq profile comparisons).* **b**) Volcano plot reporting differential gene expression levels as a function of the significance levels with a 1% alpha cutoff (light grey indicates genes with an *adjusted p* > 0.1). Respectively, red and yellow dots indicate genes that have and have not been previously associated with fruiting-body development^9–11^. The legend reports the absolute gene numbers for both categories and their frequencies relative to the total number of DE genes. For the complete list of DE genes, refer to Supp. Table S1 and Supp Table 5 for the complete list of DE genes associated with development. Visit this <u>link</u> to explore the graph in detail. **c)** Plot showing the expression-difference levels of the 42 DE genes with altered expression in the Δ*pxr* mutant during vegetative growth associated with development in all three published *M. xanthus* developmental-transcriptome studies^9–11^. Orange and blue bars highlight the increase or decrease of transcript levels, respectively. **d)** Western blot showing FruA presence/absence in the WT (GJV1) and the *Δpxr* mutant during vegetative growth (V, in red) and throughout two days of development (0, 6, 8, 12, 24 h). The arrow on the right side indicates the FruA-specific band (∼ 24.7 kDa), while the asterisk indicates the presence of an unspecific recognition of the antibody used.

### pxrA (MXAN_1079) is a likely Pxr target important for fruiting-body development in M. xanthus

The above cross-study transcriptional profile comparisons indicate that experimental idiosyncrasies can influence the identification of developmental genes. Thus, one could not exclude that other genes differentially expressed in the *Δpxr* mutant are also relevant to fruiting-body development despite not being identified as such in previous development-transcriptome analyses. This is perhaps particularly true for those gene products with an uncharacterised function^9–11,36^ (Fig. 3b and Supp. Table 1). Among those, *MXAN_1079,* encoding for a putative GNAT-acetyltransferase, was the most differentially increased transcript in the *Δpxr* mutant (Fig. 2c, 3b, and Supp. Table 1). *MXAN_1079* is predicted to be transcribed from its native promoter localised downstream of *px*r (Supp. Fig. 7a). In a previous study, the evolved strain PX bearing a non-functional *pxr* gene also exhibited increased expression of *MXAN_1079*, and this effect was hypothesised to contribute to the restored developmental proficiency of PX^12,37^. Here, *MXAN_1079* expression was consistent with the previously published data, supporting our previous hypothesis: Heightened expression of *MXAN_1079* associated with the lack of Pxr function likely allows cells to bypass developmental roadblocks. -We propose to rename *MXAN_1079* as *pxrA,* given its position next to *pxr* and its expression strongly influenced by Pxr (Supp. Fig. 7a).

To directly test whether *pxrA* is important for development, we asked whether the absence of a functional PxrA would affect fruiting-body formation. We generated a non-functional merodiploid mutant of *pxrA* in the WT genetic background (GJV1*pxrA::pCR1079*, hereafter referred to as *pxrA^−^*; see Methods: *Strains and culturing conditions*) (Supp. Fig. 7a) and monitored the developmental morphologies of *pxrA^−^* and WT populations over 141 h (ca. 6 days) by timelapse microscopy (Fig. 1b, see Methods: *Image acquisition and analysis*). Compared to WT, *pxrA^−^* cells were incapable of forming mature darkened fruiting bodies, even after 141 hours on starvation plates (Wilcoxon test relative to the end of the experiment, W = 16, *p* = 0.0294) (Fig. 4a and b). The reduced ability of the *pxrA^−^*mutant to make a lower number of mature fruiting bodies than WT was also confirmed with larger inoculation sizes (Supp. Fig. 7b). Interestingly, *pxrA^−^* cells initiated aggregating slightly later than WT cells, although this difference was non-significant with our sample size (Wilcoxon test relative to the onset of aggregate formation, W = 2, p = 0.086) (Fig. 4b). Moreover, we also couldn’t detect the second wave of development characteristic of the WT strain (Fig. 4a and b).

**Figure 4.**
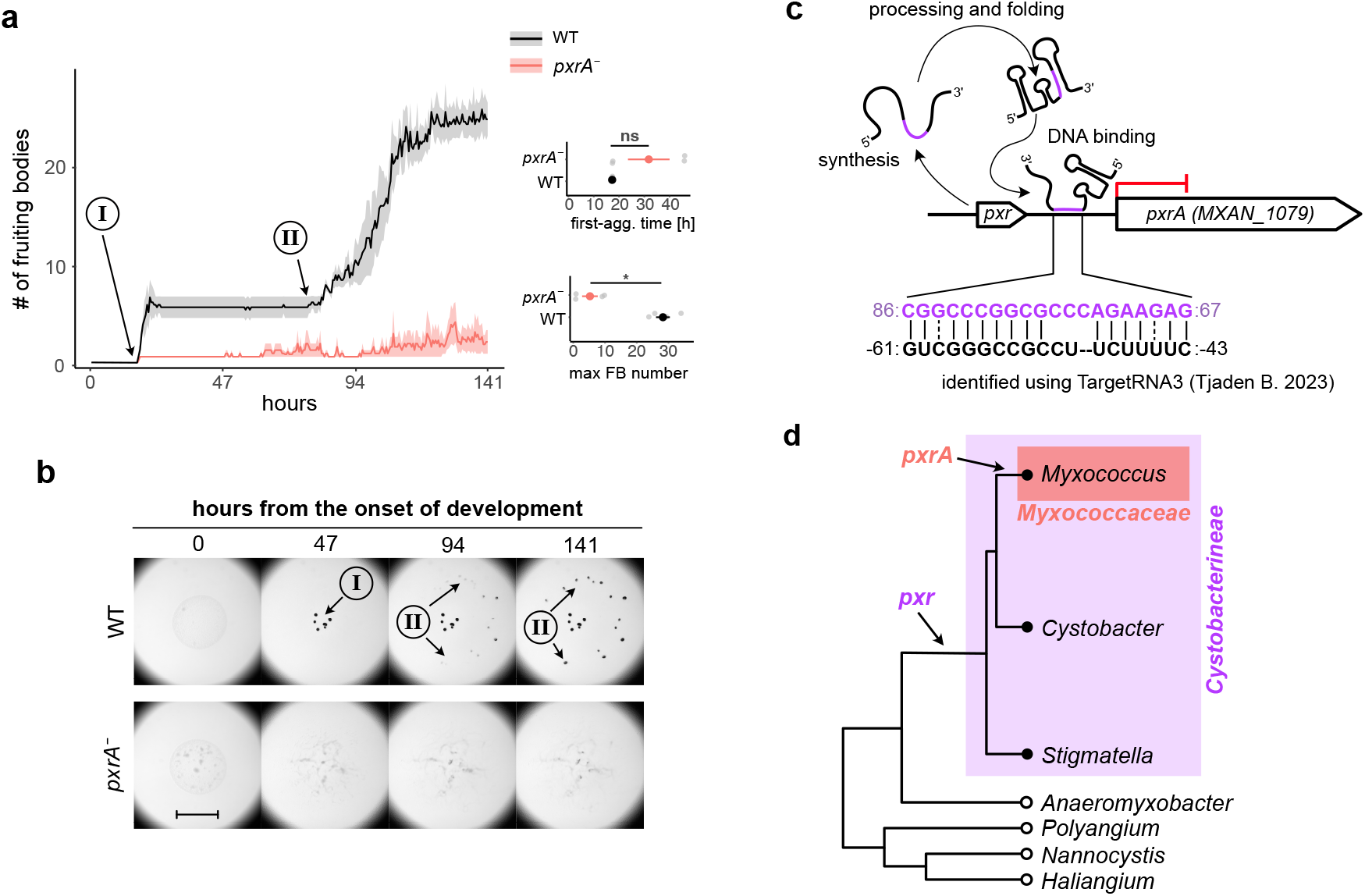
*pxrA* is a potential Pxr binding target essential to fruiting-body development. a) Plot reporting the average number of fruiting-body counts (thick lines) over time for WT (black line and shaded area) and *pxrA^−^* cells (red line and shaded area). The two developmental waves localisation of WT developing cells are indicated with I and II circled in the plot area, with arrows pointing at the corresponding time point. Graphs on the right report the average developmental time for the first aggregates to form (top) and the average number of fruiting bodies at the end of the experiment (bottom) for WT and *pxrA^−^* coloured in black and red, respectively. The shaded areas in the large plot and the error bars in both graphs represent the standard errors associated with each measurement (n = 4). The asterisk indicates a significant mean difference, while *ns* indicates a non-significant one. **b**) Representative images of developing fruiting bodies (dark spot) over the time course for WT (GJV1) (top row) and *pxrA^−^* cells (bottom row). The circled-ordinal numbers I and II and arrows indicate the emergence and localisation of the two waves observed in the WT. The scale bar equals 1 mm. **c)** Schematic diagram showing the effect and the potential binding site of Pxr to the upstream promoter region proximal to *pxrA (MXAN_1079).* Numbers at the end of each string of letters indicate the base position relative to the Pxr sequence (purple numbers and letters) and the *pxrA* transcriptional start site (black number and letters). For a complete list of all predicted Pxr binding targets, refer to Supp. Table 6. **d**) Phylogenetic tree qualitatively contrasting the emergence of *pxr* and *pxrA* in myxobacteria (modified from ref. ^15^ with individual species collapsed to individual genera). Refer to ref ^15^ and Supp. Fig. 6c for a more detailed visualisation of pxr and PxrA phylogenies, respectively.

In addition to experimentally demonstrating that *pxrA* plays a significant role in positively regulating the *M. xanthus* developmental program, we also sought to bioinformatically identify potential Pxr targets by using TargetRNA3^38^ to run an unbiased sequence alignment. The resulting list of 25 potential Pxr binding regions included a sequence near the predicted promoter region of *pxrA* (Supp. Table 6). The portion of the Pxr sequence predicted to bind upstream of the *pxrA* promoter is located primarily within the third of three predicted Pxr stem-loop structures known to be necessary for Pxr function^39^ (Fig. 4c).

Surprisingly, none of the other 24 predicted binding regions was associated with a gene differentially expressed in *Δpxr*. Yet, the effect of Pxr on these transcripts might affect translation without altering the transcript abundance. Moreover, none of these potential targets showed significant similarities in their functional annotations.

We then asked whether the phylogenies of PxrA and Pxr might be similar given their regulatory interaction (see Methods: *Phylogenetic analysis of PxrA)*^15^. Our analysis showed homologous PxrA sequences between a few myxobacteria species (*M. macrosporus, M. stipitatus, M. virescens,* and *M. xanthus,* among other non-characterised *Myxococcus* strains), all of which belong to the suborder Cystobacterineae within the family Myxococcaceae, where *pxr* is thought to have emerged (Fig. 4d, Supp. Fig. 7c)^15^. Thus, *PxrA* emerged most likely later than *pxr* and was retained or transferred into only a relatively small number of species, despite its current critical role in *M. xanthus* aggregative multicellularity (Fig. 4d).

Altogether, our results identify *pxrA* as a novel positive regulator of fruiting-body development and a likely direct target of Pxr regulation.

## DISCUSSION

In bacteria, sRNAs regulate gene expression and are arguably associated with all major biological processes^1–3^. Pxr is a *trans*-acting sRNA involved in fruiting-body formation in myxobacteria, an aggregative multicellular process that takes place in response to starvation. Specifically, Pxr prevents the activation of the developmental program at nutrient levels abundant enough to support extensive vegetative growth^5^. Here, we have expanded our understanding of the phenotypic effects of Pxr regulation by showing that this sRNA not only prevents development under high-nutrient conditions, but also limits the pace of development under starvation conditions. We further identified genes regulated by Pxr during vegetative growth by comparing the transcriptomic profiles of cells lacking or expressing *pxr*, including the key developmental regulator *fruA* and a GNAT-acetyltransferase gene immediately downstream of *pxr – pxrA –* that we reveal as necessary for fruiting-body formation. We additionally identified many possible direct binding targets of Pxr, including the promoter region of *pxrA*.

### The role of pxr in controlling the dynamics of development

We showed that the absence of *pxr* expression greatly accelerates both the initiation and completion of development when nutrients are absent (Fig. 1). Until now, Pxr has been understood to control the nutrient conditions under which development is initiated; our new results show it also is a major regulator of developmental timing. Interestingly, the observation that the WT and the *pxr* mutant had similar numbers of fruiting bodies at the end of the experiment indicates that the absence of *pxr* expression affects only the temporal dynamics of development without interfering with the developmental potential of cells. We need more information about the molecular details of how Pxr function can alter developmental timing. Our data suggest a potential role of Pxr both before and after the initiation of development. For example, it is plausible that the unprocessed Pxr isoforms are involved during development (i.e., Pxr-L and Pxr-XL), which, in contrast to the cleaved Pxr-S, remain abundant throughout the aggregative process^5,13^.

The absence of *pxr* expression can impact the dynamics and total duration of development beyond the induction of aggregative multicellularity. Thus, variation in *pxr* expression across different genotypes could have profound implications for myxobacteria ecology and evolution. For example, mutations that lead to an early commitment to the developmental program while nutrients can still favour growth could negatively affect performance at interference competition mechanisms requiring high metabolic activity^40,41^. Alternatively, being able to induce sporulation earlier and more quickly could be advantageous in resisting such antagonistic behaviours exerted by co-developing genotypes that develop more slowly^41,42^. Our data brings important evidence suggesting an ecological role of *pxr* exerted by controlling the developmental dynamics of multicellular development.

### Lack of pxr expression influences the transcriptional levels of metabolic genes

We found that deletion of *pxr* affected the transcript levels of 4% of all annotated *M. xanthus* genes (299/7451 total DK1622 annotated genes). However, it is likely that Pxr directly interacts with only a small minority of them, with the majority being regulated indirectly. For example, the congruency in the sign of transcriptional effects within operons supports this hypothesis.

More than half of the genes affected by the absence of *pxr* have been linked to metabolism (Fig. 2). Notably, association with metabolism does not exclude potential involvement in development. Interestingly, metabolism-related genes had decreased transcript values in cells without *pxr* expression. By reducing metabolic gene transcripts, we hypothesise that the lack of Pxr pried cells toward a developmental state where metabolic processes are typically turned off or reduced (Fig. 5).

**Figure 5.**
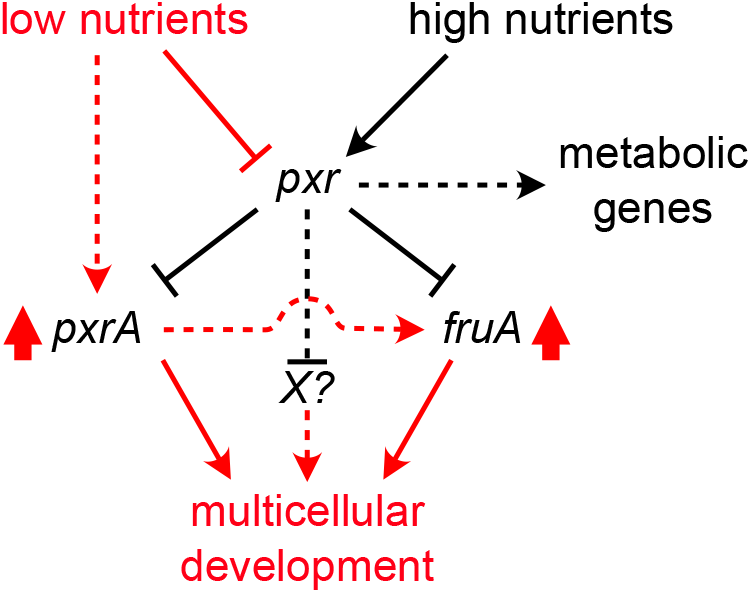
Hypothetical model of Pxr function preventing multicellular development by repressing levels – and hence activity – of key regulators of fruiting-body development. When nutrients are conducive to growth (black lines and arrows), Pxr represses levels of positive developmental regulators such as PxrA and FruA. Moreover, under the same conditions, our data suggest a positive effect of Pxr on the expression of genes associated with metabolic processes. When nutrient levels become scarce (red lines and arrows), however, the repressive function of Pxr is relieved. Cells decrease their metabolic rates and activate the developmental program via *pxr* repression, leading to multicellularity by increasing the expression of positive regulators of fruiting-body development.

### Altered developmental gene expression in the absence of Pxr

In line with our initial expectation, we also strengthened the association between *pxr* and multicellular development from a molecular standpoint. Using previously published lists of potential *M. xanthus* developmental genes^9–11^, we determined that more than half (67.6%) of the genes differentially expressed in the *pxr* mutant were associated with development in at least one of three developmental- transcriptome studies (Fig. 3). We found a number of critical developmental genes, such as *nsd, MXAN_0736, pxrA* and *fruA* (among others) showed transcriptional levels consistent with the expectation of an activated developmental program in cells lacking *pxr* expression.

During fruiting-body morphogenesis, FruA, a DNA binding response regulator, is essential in orchestrating population rippling, aggregation, and sporulation^11,27^. Transcription of *fruA,* depending on the early developmental A-signal, is induced after 3-6 hours of starvation in WT cells. The function of the FruA protein controlled by the contact-dependent C-signal was thought to be activated by phosphorylation. However, the cognate histidine kinase remains unknown^43^. Here, we showed that the increase of *fruA* transcript was indeed followed by an increase of the FruA protein levels detectable during vegetative growth. Our data suggests that the accelerated development observed for *Δpxr* cells may directly result from the differential expression of developmental genes already during vegetative growth (Fig. 5).

We also found developmental genes important to development less abundant during vegetative growth in the *pxr* mutant cells than in WT. For example, the two developmental genes *lonD* (also known as *bsgA,* important for the early developmental signally pathway) and the response regulator *rodK,* whose functions are relevant to fruiting-body formation^29,31,35^, had decreased transcript levels in *pxr* mutant cells (Supp. Table 1 and 5). However, in the case of *lonD*, it has been indicated that a *pxr* mutant can undergo development by bypassing the requirement of the *lonD* gene, indicating that the removal of the developmental gatekeeper Pxr can trigger a developmental process by turning on suites of downstream developmental regulons without a functional *lonD* gene^13^. Therefore, high expression of *lonD* is not required in a ready-to-develop state of the *Δpxr* mutant cells. It is possible that a reduction in *lonD* transcripts is a byproduct of an auto-regulatory circuit to shut down early signalling genes that are not needed.^64^

The obtained lists of developmental genes differ between the three consulted studied (Supp. Fig. 4a)^9–11^, and the temporal dynamics and expression levels of those genes shared between the studies may vary, too. These differences are likely due to protocol differences and genetic backgrounds (different reference strains), but study idiosyncrasies may also reflect the molecular complexity and plasticity intrinsic to fruiting-body development^11,44–46^. Discrepancies in specific gene-expression levels from our transcriptome profiles can be ascribed to the different physiological states in which cells were sampled (growing vs starving cells).

### pxrA is a potential Pxr binding target associated with multicellular development

In myxobacteria, the functions of many genes remain uncharacterised, and in many cases, function is predicted only based on sequence similarity. Indeed, one of the clusters of orthologous genes enriched for genes differentially expressed in the *pxr* deletion-mutant (cluster S) includes genes of unknown function, including *MXAN_1079,* which encodes a putative GNAT-acetyltransferase^37^. GNAT- acetyltransferases are conserved in all organisms and are involved in post-translational modifications that modulate many cellular functions^47^. The transfer of an acetyl group from acetyl CoA to a protein substrate can occur at either the amino-terminal end or at the ε-amino group of an internal lysine residue. Post-translational modification by acetylation is considered a major mode of regulating gene expression and is as prominent as phosphorylation-mediated modification.

The simple explanation for the increased *pxrA* transcripts detected in a *pxr*-deletion mutant is that *pxrA* is a binding target of Pxr that is negatively controlled by this sRNA. Indeed, bioinformatic analysis does predict base-pairing between Pxr and the *pxrA* non-coding region upstream to its transcriptional start site. However, the possibility that the upstream localisation of the *pxr* deletion might have contributed to the specific transcriptional levels of this gene cannot be excluded. Yet, the upregulation of *pxrA* in response to an impaired function of *pxr* reflects previous observations done with a different genetic background to this study, where the association of the Pxr-*pxrA* interaction with development was first hypothesised^12^.

Here, we went one step further and demonstrated that, besides being a potential Pxr target, *pxrA* is critical to fruiting-body formation and developmental timing (Fig. 4). Interestingly, our phylogenetic analysis indicated that *pxrA* emerged in the Myxobacterineae family, a subgroup of the Cystobacterineae in which *pxr* is considered to have originated^15^ (Fig. 4). Our analysis suggests that *pxrA* emerged more recently than *pxr*. It will be interesting for further work to investigate potential developmental features unique to the interaction between these two genes in species that carry them both.

A previous study found another putative GNAT-acetyltransferase (*MXAN_6704*) important to fruiting body-formation^48^. As for *pxrA,* none of the consulted developmental transcriptome studies^9–11^ had associated *MXAN_6704* with development. In our research, *pxrA* was the only acetyltransferase to show increased transcriptional levels. Thus, these two acetyltransferases are likely regulated independently via different signalling pathways during fruiting-body development. GNAT-acetyltransferase activity is critical to gene regulation in bacteria^47^, making it tempting to speculate that the genetic interaction between *pxr* and *pxrA* is important for modulating developmental gene expression. A further analysis contrasting the molecular targets of both PxrA and MXAN_6704 could help understand the role of acetyltransferases during bacterial aggregative development.

Our analysis indicates that classifying a gene as *developmental* depends, to some degree, on the environmental context, the experimental procedure used, and the intrinsic plasticity and molecular redundancy of the fruiting-body formation process^11,45^. Our demonstration that *pxrA (MXAN_1079)* positively regulates development despite the other transcriptome studies not having flagged this gene is a clear example of this scenario, which suggests that additional new developmental genes remain to be discovered.

Future research themes of interest include the exact molecular mechanisms used by Pxr to exert its function, experimental confirmation of Pxr binding targets, and what role *pxr* expression plays in the evolution of the temporal dynamics of aggregative multicellular development across the myxobacteria.

## METHODS

### Strains, culturing conditions and induction of development

The following strains were used for this study: GJV1 (a laboratory-derived strain of DK1622^49^ used as wild-type reference (WT); GJV1*Δpxr* (a *pxr*-deletion strain^5^) lacking the expression of the *pxr* gene (referred to as *Δpxr* in the main text); and GJV1*pxrA*::pCR-1079 (a strain characterised by the knockout of *pxrA* (*MXAN_1079*), referred as *pxrA^−^* in the text; see Methods: *Plasmid and strain construction*). Myxobacteria strains were grown in the casitone-based liquid media CTT (Tris-HCl 10mM, MgSO4 8mM, Bacto Casitone 1 %, KH2PO4-K2HPO 1mM, pH 7.6). Bacterial cultures were incubated at 32 °C with orbital shaking at 300 rpm. To induce fruiting-body formation, cells were plated on nutrient-limited TPM agar plates (Tris-HCl 10mM, MgSO4 8mM, Bacto agar 1.5 %, KH2PO4-K2HPO 1mM, pH 7.6). For developmental assays, mid-log cultures were pelleted and resuspended in TPM liquid buffer (Tris-HCl 10mM, MgSO4 8mM, KH2PO4-K2HPO 1mM, pH 7.6) to ∼5 x 10^9^ cells/mL, after which 50 µl or 1 µl (for high-magnification developmental analysis under Nikon Ti2 microscope) samples were spotted on a TPM-agar plate.

### Plasmid and pxrA knock-out strain construction

To make the pCR-1079 plasmid, an ∼280-bp internal fragment of the *pxrA* coding gene was generated by a PCR reaction with primers 1079-11 (TCTTTGCCCGGCTGTTTCTTG) and 1079-12 (CCCGCTTCACGTTGAGGAC) and cloned into the pCR-Blunt vector. Restriction-enzyme digestion and Sanger sequencing confirmed the positive constructs. The *pxrA-*knockout strain GJV1*pxrA*::pCR-1079 (referred to as *pxrA^−^* in the main text) was constructed by transforming the parental strain GJV1 with pCR-1079 and isolated from a kanamycin- CTT hard agar plate. Integration of pCR-1079 at the *MXAN_1079* locus created a non-functional merodiploid with one truncated *MXAN_1079* copy lacking codons for residues 113-278 of the encoded protein’s C-terminus and the other partial copy lacking codons for residues 1-19 of the protein’s N-terminus (see Supp. Fig. 7a). The construct of the merodiploid mutant was verified with Sanger sequencing using the same primers indicated above and vector-specific primers.

### RNA extraction and RNA-seq analyses

RNA isolation from three independent biological replicates of the strains GJV1 and GJV1*Δpxr* was done using the RNeasy Mini Kit (QIAGEN, # 74004) according to the manufacturer’s protocol using cells at mid-log phase. The isolated RNA was treated with the TURBO DNA-*free*^TM^ Kit (Invitrogen, # AM1907) in a volume of 50 µl using 5 µg of total RNA, and the reaction was incubated at 37 °C for 45 min. Ribosomal RNA (rRNA) was depleted using Ribo-Zero Plus rRNA Depletion Kit (Illumina, #20040526) according to the manufacturer’s protocol. RNA quality was checked using the Aligent 4150 TapeStation® machine, and only samples with an RNA integrity number (RIN) higher than 8 were further processed. The Illumina True-seq protocol for total RNA was used for library preparation, and 150bp single-end read sequencing was run on an Illumina-Novaseq 6000 at the Functional Genomic Centre Zurich (FGCZ), University of Zurich, Switzerland (Supp. Fig. 2b). Sequencing reads were checked for quality using *FastQC* v0.73^50^ and trimmed with *Trimmomatic*^51^ for the adapter sequences and poorly sequenced bases (parameters set at 4:20). Trimmed reads were then mapped to the DK1622 (ASM1268v1) genome using *bowtie*^52^ while controlling for correct strand orientation. Read counts per gene were obtained using *HTSeq*^53^ with minimum alignment quality set to 10 bases and model *union* to obtain the final read counts. On average, the number of mapped read counts per individual sample was 2 million. To assess differences in gene expression levels, read counts per gene of all six independent replicate samples (three replicates for GJV1 (WT) clones and three replicates for GJV1*Δpxr* clones) were analysed with *DESeq2*^54^ using the quantile method to normalise read counts and compared across replicates. Significant differences were considered using an *α* value set at 0.1. RNA-seq-specific analyses were all conducted on the Galaxy platform (https://galaxyproject.org)^55^. Enrichment analyses for the clusters of orthologous genes (COGs) and operons were performed with *FUNAGE-Pro*^56^ using the DK1622 (ASM1268v1) as the reference genome. Networks of gene associations and GO enrichment analysis were done using the STRING database^18^ with the *stringApp* on the Cytoscape software^57^ with the genome DK1622 (ASM1268v1) as reference. In addition, we retrieve the individual gene functional annotations from the UniProt^58^ database and store them as Supp. Data 1. The raw *fastq* files and read counts per gene are deposited at GEO with the following IDs (GSE265958). We generated an online application at the following link (https://jzego7-marco.shinyapps.io/rnaseq_app/) with Shiny^59^ (https://shiny.posit.co/) in R^60^ to allow a user-friendly exploration of our data. The application is stored as Supp. Code listed among the Supplementary Information.

### Comparisons of RNA-seq profiles

A list of *M. xanthus* genes associated with fruiting-body development was obtained from three studies that used RNA-seq to characterise transcriptional profiles of cells during development^9–11^. The degree of similarity between our study and the others was assessed by comparing the expression levels of cells sampled during the exponential growth phase. Each study differed in some respects (e.g., growth media, equipment used for growth), with the vegetative growth of wild-type cells being the only experimental condition common to all four studies. When available, expression levels were recovered as normalised gene counts^911^ or as the differential expression analysis^10^. However, all transcriptional levels were scaled across their relative mean value before direct comparison. Pearson’s correlation test was performed for all pairwise combinations across all datasets, including ours (Supp. Fig. 5a). As mentioned, the three studies differ in their experimental setups (e.g., the protocol used to process and sequence the RNA and the genetic background of the reference WT strains). Therefore, significant differences between them were expected, as previously acknowledged^10,11^. Given these considerations, a correlation close to 50% was heuristically considered relevant. Thus, all three other studies were used to extract potential developmental genes.

### Image acquisition and analysis

Analyses of fruiting-body development over time for the two strains GJV1 and GJV1*Δpxr* were done with a Nikon Ti2-E microscope and a DS-Qi2 camera. Developing cells were observed for a maximum of 141 h, and images were taken every 30 m with a 2x objective. While at the microscope, cells were kept at 32 °C using a customised microscope stage-top incubator. Development time lapses were analysed with ImageJ software v2.9.0^61^ to assess the number of fruiting bodies over time (Supp. Data 2). Using a previously established protocol in our lab^62^, fruiting bodies were identified based on their grey values and consistently counted across all samples. Subsequently, statistical analysis of the data obtained was run in R v4.0^60^.

To obtain representative images of fruiting bodies on plates after 5 days of starvation-induced development, a Zeiss STEMI 2000 microscope and a Nikon Coolpix S10 camera were used.

### Phylogenetic analysis of PxrA

PxrA (MXAN_1079) orthologs were identified by searching all sequences in the non-redundant NCBI and integrated microbial genomes (IMG) databases^63^. Next, hits for E-values lower than 10^-100^ and sequences higher than 80% similarity were selected and aligned against each other using MUSCLE^64^. ProtTest 3 ^65^ was used to determine the optimal model (JTT+G+F, with gamma: 0.67) before the tree generation. Maximum Likelihood trees were generated using PhyML 3.0 ^66^, running 1,000 bootstraps. Lastly, iTOL v4 ^67^ was used to visualise the tree obtained.

### Western immunoblotting assay of FruA

The protein samples prepared from the vegetative cultures growing in CTT (V) as well as developing populations submersed in MC7 buffer at different time points (0, 6, 12, 18 and 24 h) were electrophoresed in a 12.5% SDS polyacrylamide gel. Subsequently, proteins were electro-transferred from electrophoresis gels to Whatman Protran nitrocellulose membranes by a semidry blotting device. The membrane blots were first hybridised with rabbit anti-FruA antibody (1:1500 dilution in TTBS [Tris-buffered saline with 0.1% Tween 20] -1% gelatin) followed by incubating with alkaline phosphatase (AP)-conjugated goat anti-rabbit immunoglobulin G (1:3000 dilution in TTBS-1% gelatin). Detection of FruA protein was visualised for all conditions in parallel by a chemiluminescent reaction with a light emitting substrate (CDP- Star™) (Ambion).

## ACKNOWLEDGEMENTS

We thank the Genetic Diversity Centre (GDC – ETH Zürich), the Functional Genomic Center Zürich (FGCZ – University of Zürich) for the helpful technical support and the Evolutionary Biology group at ETH Zurich for comments on the manuscript. This work was partly funded by an EMBO Long-Term Fellowship (ALTF 1208–2017) to **M.L.F**. and Swiss National Science Foundation grant 310030B_182830 to **G.J.V.**

## AUTHORS’ CONTRIBUTION

**Y.-T.N.Y., G.J.V. and M.L.F.** designed the project, experiments and analyses; **M.L.F.**, **J.V.**, **S.C.**, **S.E.,** and **Y.-T.N.Y.** conducted the experiments; **M.L.F.** and **J.V.** analysed the data and generated figures. **M.L.F.** and **Y.-T.N.Y.** drafted the manuscript; all authors reviewed and commented on the manuscript; **M.L.F., G.J.V.** and **Y.-T.N.Y.** revised the manuscript.

## DATA AVAILABILITY

RNA-seq data are deposited at NCBI’s Gene Expression Omnibus^68^ and are accessible through GEO Series accession number GSE265958 (ncbi.nlm.nih.gov/geo/query/acc.cgi?acc=GSE265958).

Supplementary Information is stored on Dryad at https://doi.org/10.5061/dryad.xsj3tx9pn

## Supplementary information

**Supp. Fig. 1.**
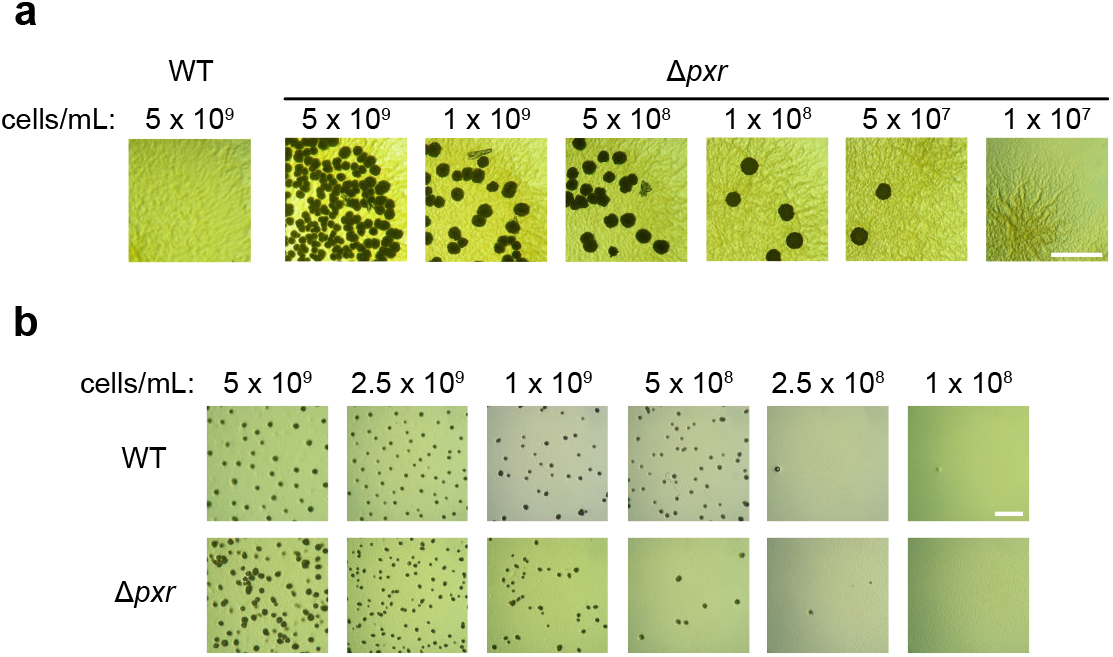
**a**) Microscopy images comparing five-day-old WT and *Δpxr* cell cultures inoculated on an agar plate with 0.3% casitone, with decreasing cellular densities for the *Δpxr* mutant. Dark spots are mature fruiting bodies. **b**) Microscopy images showing five-day-old cultures of WT and *Δpxr* (top and bottom array rows, respectively) after inoculation onto buffered (TPM) agar at six different initial densities. Dark spots are mature fruiting bodies. Scale bars equal to 2mm for both panels.

**Supp. Fig. 2.**
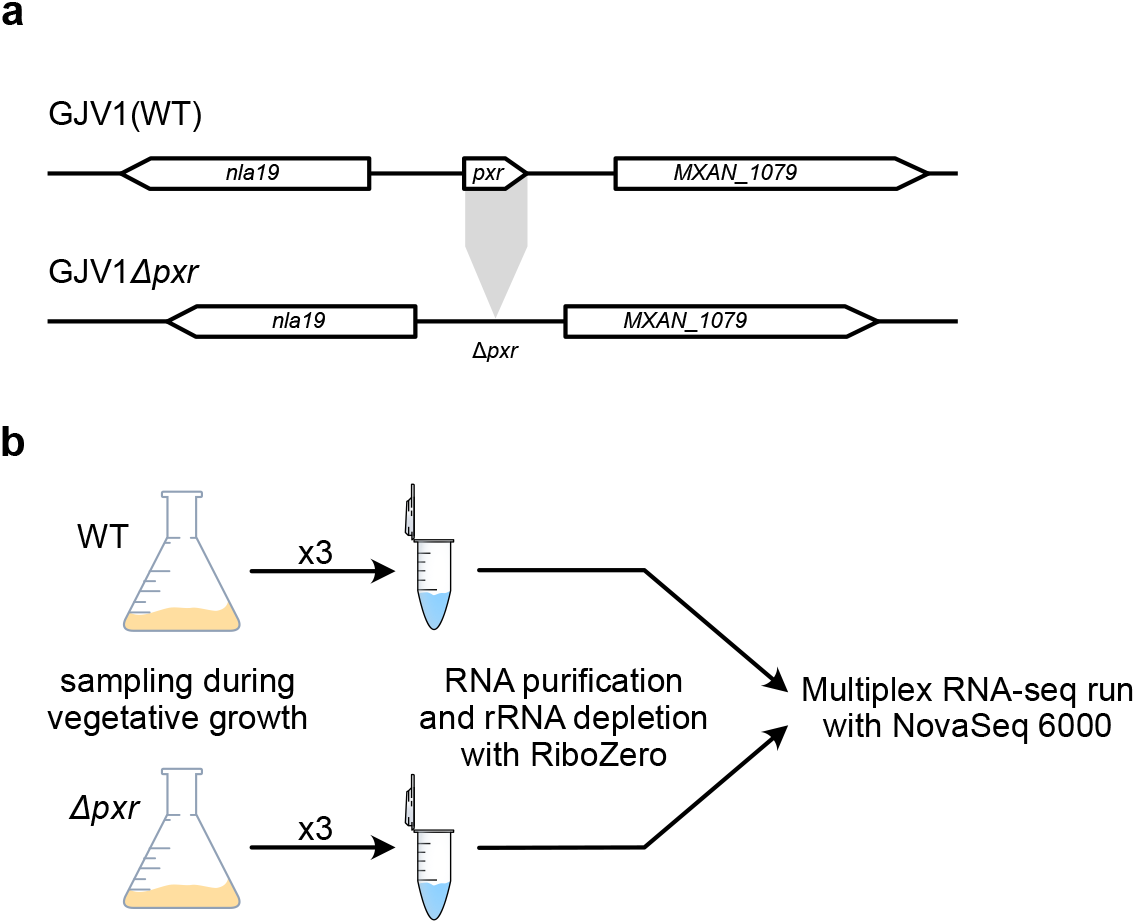
**a**) Schematic representation of the *pxr* locus in WT (GJV1) cells (top) and GJV1*Δpxr* mutant cells (bottom). **b)** Cartoon reporting the experimental design used to sample RNA from cells during vegetative growth in rich media. RNA was extracted, purified, and rRNA removed using RiboZero before whole transcriptome sequencing on a NovaSeq 6000 machine. The total number of reads per sample was estimated to be approximately eight million (total number of annotated genes for DK1622 = 7579).

**Supp. Fig. 3.**
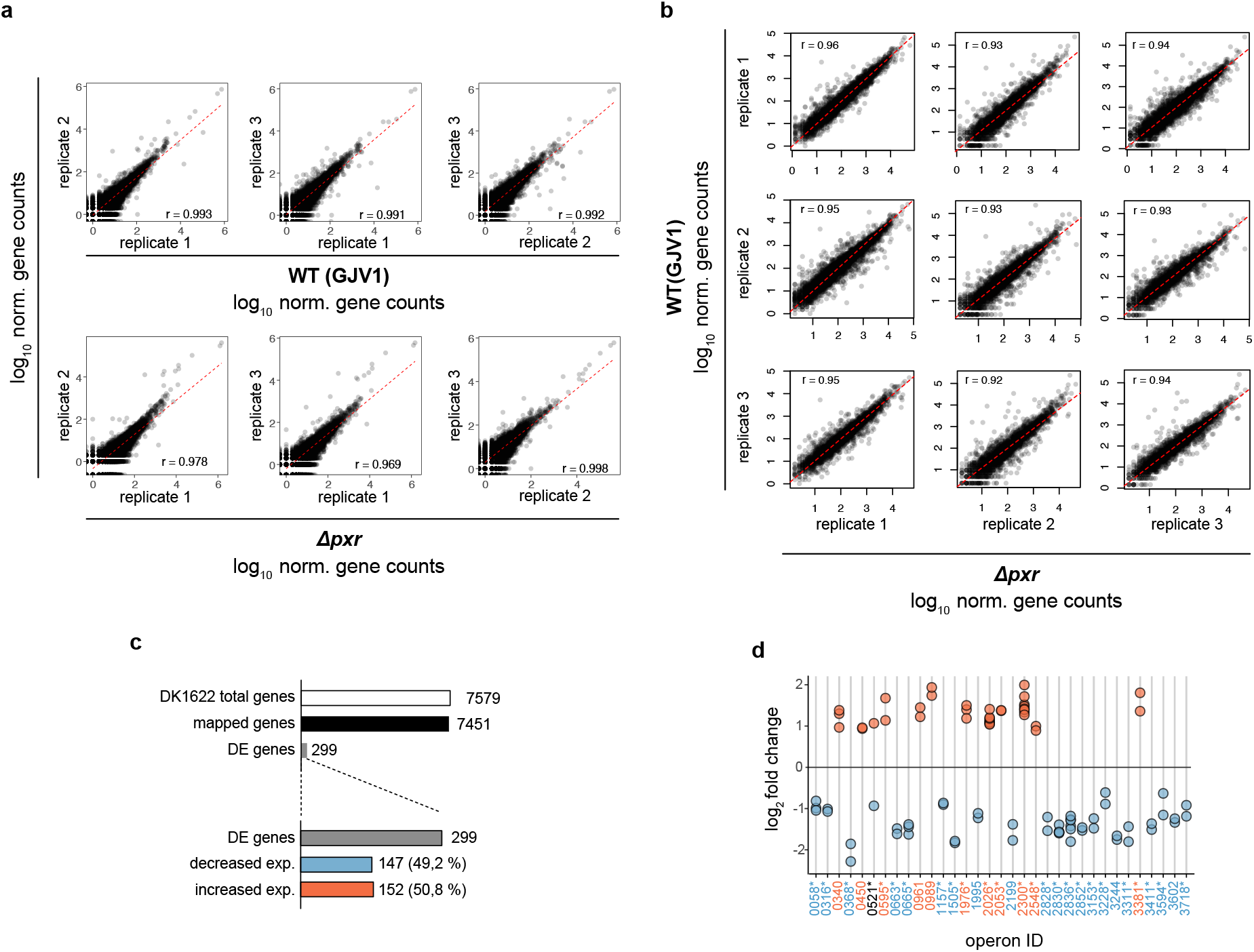
**a**) Scatter plots reporting the pairwise comparison of gene counts between the individual biological replicates per each strain (WT or *Δpxr*). Pearson’s coefficients of correlation (*r*) are reported within each plot. We measured an average variance value of 0.048 and 0.142 for WT and *Δpxr,* respectively. **b)** Scatter plots reporting the pairwise comparison of gene counts between WT (GJV1) and *Δpxr* individual biological replicates. Pearson’s coefficients of correlation (*r*) are reported within each plot. **c)** Top: Summary of the total number of unique transcript sequences mapped onto the DK1622 reference genome (white and black bars, respectively), the number of genes differentially expressed (DE) between WT and the Δ*pxr* mutant (grey bars) compared to the mapped and total genes. Bottom: Numbers (and percentages) of DE genes (grey bar) that are down-regulated (blue bar) and up-regulated (red bar). **d)** Dot plot illustrating the level of differential expression of genes (*y* axis) belonging to the list of the enriched operons (*x* axis). The blue and red text refers to operons containing genes with decreased and increased transcript levels. *0521* (in black) is the only operon that contains genes with different signs of expression levels. Asterisks placed next to the operon IDs indicate operons that contain at least one gene associated with development (75% of the 33 operons); see Supp. Table 2 and 5 for more details.

**Supp. Fig. 4.**
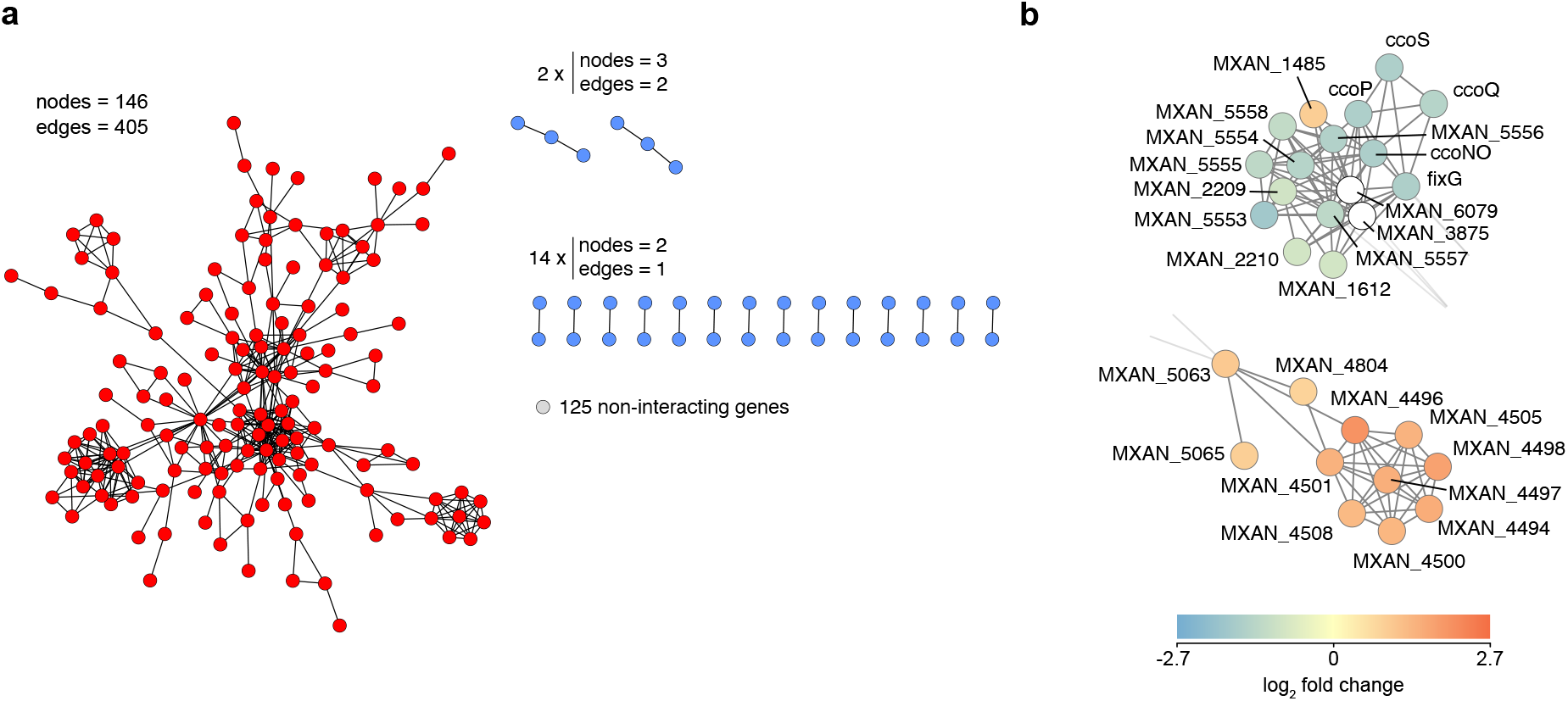
**a**) Summary of all the networks resulting from previously annotated gene interactions. The main network is red, blue which all interactions are less than five genes, and grey, which are all single non-interacting genes. **b)** Close-ups of two dense sub-clusters from the main red network reported in (**a**). Nodes are coloured by their differential expression levels, going from blue (low transcript levels) to red (high transcript levels). The depicted networks can be downloaded and consulted in greater detail online at this **<u>link</u>**.

**Supp. Fig. 5.**
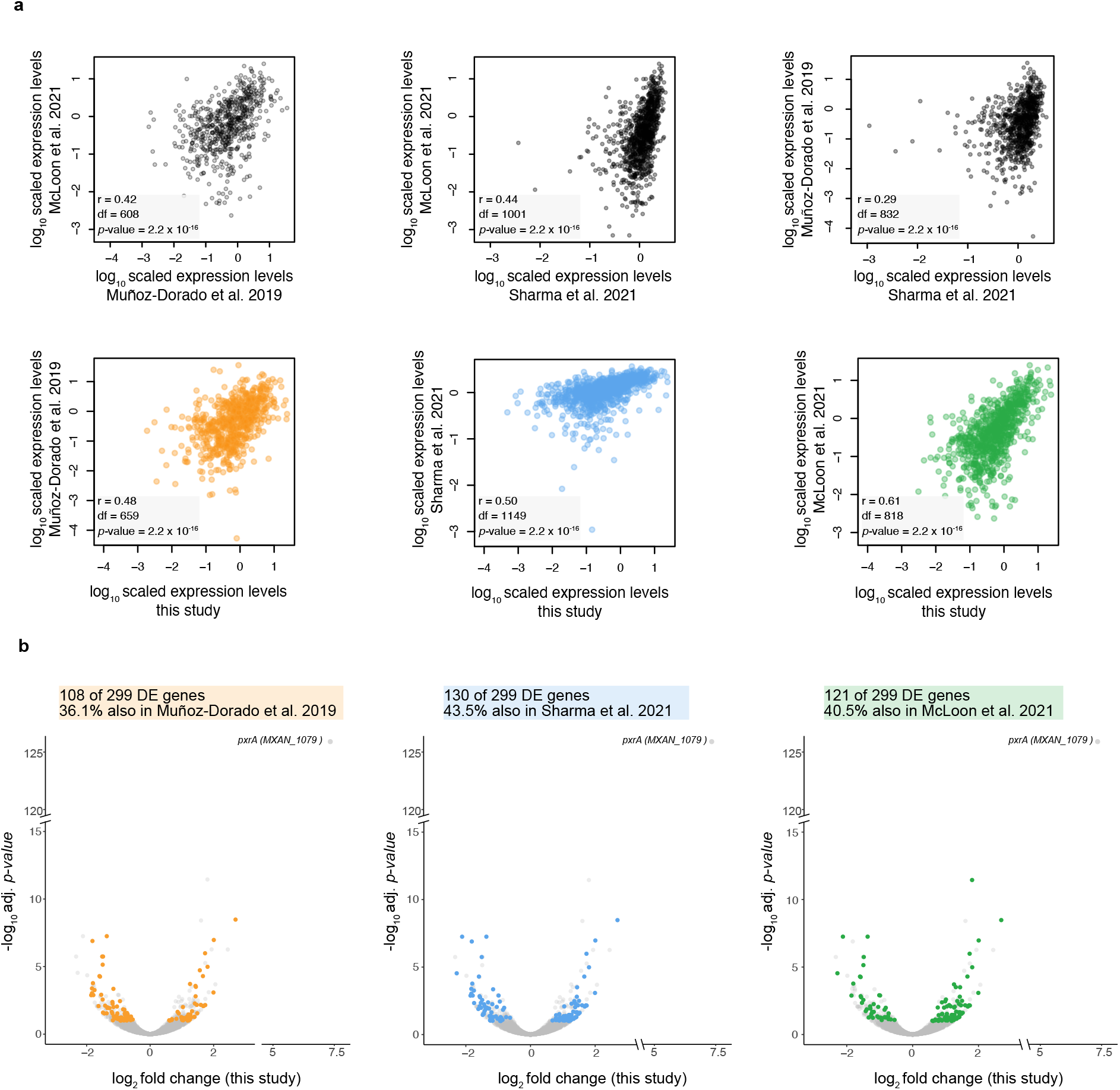
**a**) Top row: three scatter plots with pair-wise contrasts of transcript levels from vegetative cells across the three studies we used to identify developmental genes. Bottom row: three scatter plots contrast the transcript levels of wild-type vegetative cells obtained in our study vs Muñoz-Dorado *et al.* 2019 (orange dots), Sharma *et al.* 2021 (blue dots) and McLoon *et al.* 2021 (green dots). In all scatter plots (top and bottom rows), the summary statistic of the Pearson correlation test (*r*) are reported within each graph (df = degrees of freedom). **b)** Volcano plots reporting differentially expressed (DE) genes found in our study that were previously associated with development by Muñoz-Dorado et al. 2019 (orange dots), Sharma et al. 2021 (blue dots) and/or McLoon et al. 2021 (green dots). A summary of statistics for each comparison is reported above each graph. For more details on comparing RNA-seq profiles, consult Methods*: Comparisons of RNA-seq profiles.* Visit the following <u>link</u> to explore the RNA-seq data in more detail.

**Supp. Fig. 6.**
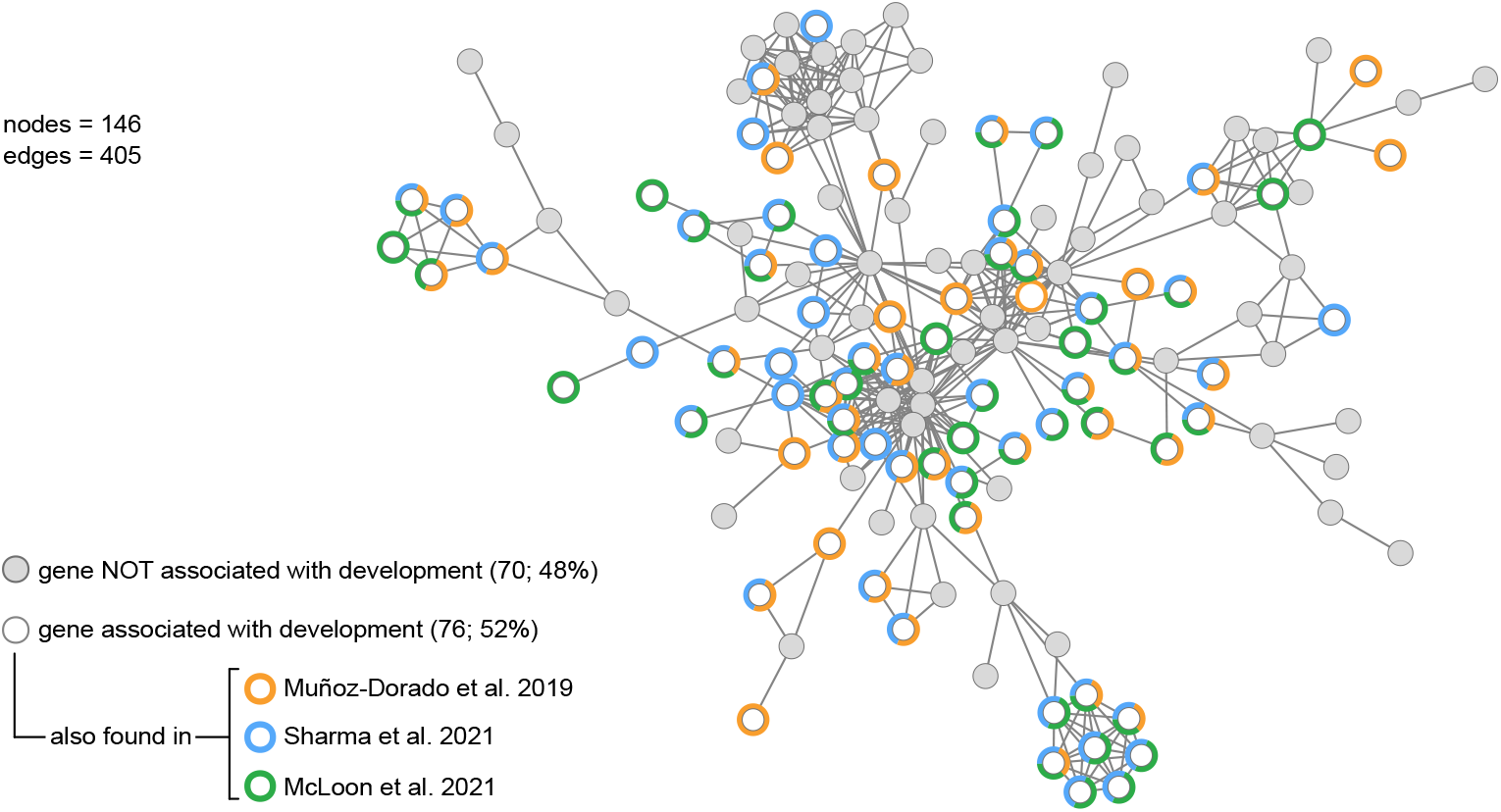
**a**)The main gene interaction network (also in Fig. 2d- and S4) highlights developmental genes in white and genes not associated with development in grey. Coloured circles identify developmental genes present in Muñoz-Dorado et al. 2019 (orange circles), Sharma et al. 2021 (blue circles), and/or McLoon et al. 2021 (green circles). The depicted network can be downloaded and visualised in greater detail using this **<u>link</u>**.

**Supp. Fig. 7.**
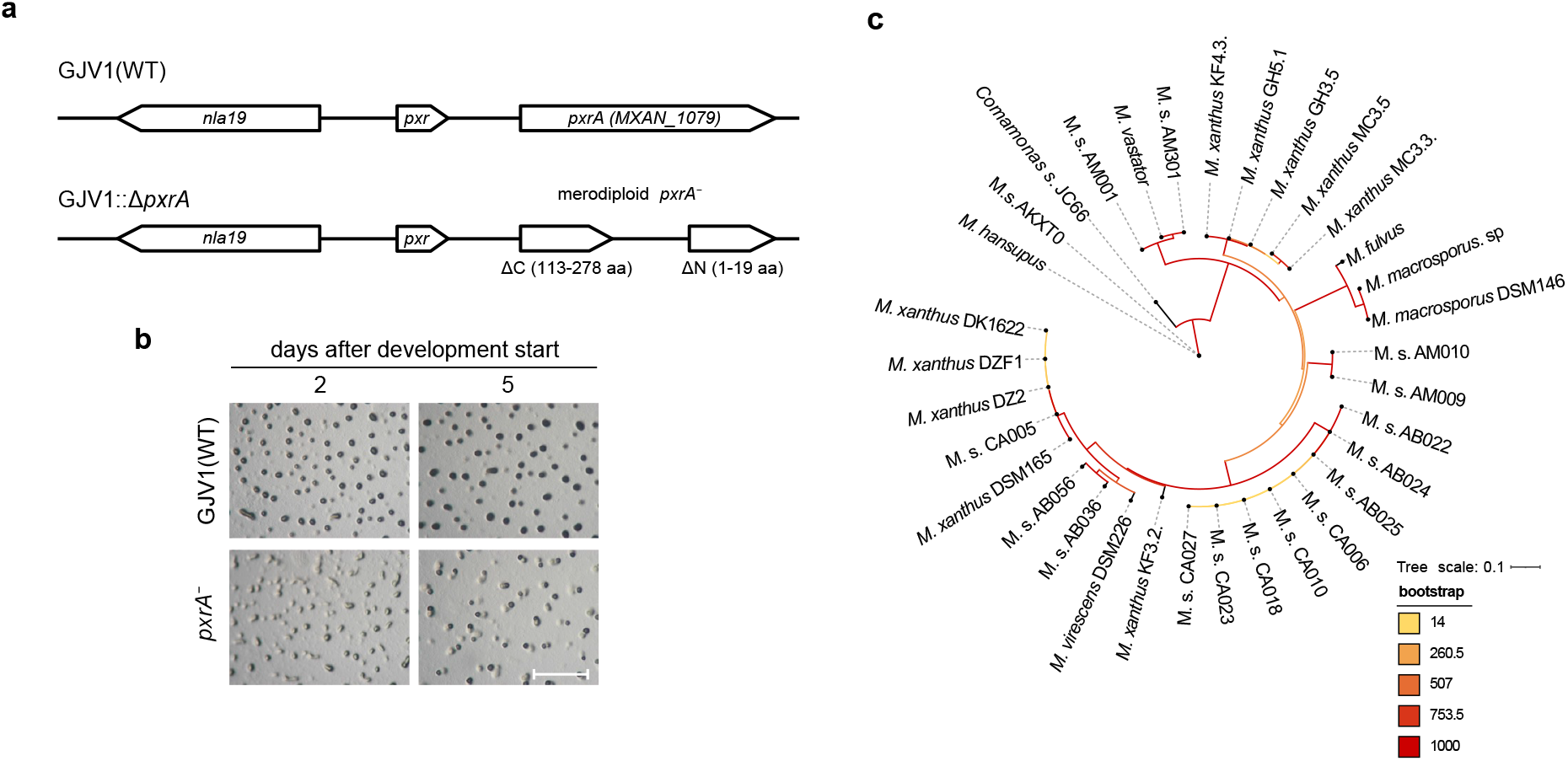
**a**) Illustration of the *pxrA* locus in WT(GJV1) and *pxrA^−^* (GJV1*pxrA::pCR-1079*) mutant cells. Merodiploid mutant cells carried the two deletions at the N- (1-19) and C-terminal (113-278) regions of PxrA (total protein length 278 aa). **b)** Representative microscopy images showing the wild-type GJV1 (top row) and the mutant *pxrA^−^* during starvation- induced development. The photographs were taken two and five days after the onset of development. The fruiting bodies are visible as darkened aggregates. The scale bar equals 1 mm. **c)** Phylogenetic tree for PxrA orthologues. Branches are coloured by their bootstrap values, and their lengths indicate the expected number of substitutions per site. The species name and strain are indicated at the end of each branch. M.s. = *Myxococcus sp*.

